# Inflammasome- and gasdermin D-independent IL-1β production mobilizes neutrophils to inhibit antitumor immunity

**DOI:** 10.1101/2020.08.04.235796

**Authors:** Máté Kiss, Lieselotte Vande Walle, Els Lebegge, Helena Van Damme, Aleksandar Murgaski, Junbin Qian, Manuel Ehling, Samantha Pretto, Evangelia Bolli, Jiri Keirsse, Yvon Elkrim, Maria Solange Martins, Amelie Fossoul, Diether Lambrechts, Massimiliano Mazzone, Andy Wullaert, Mohamed Lamkanfi, Jo A. Van Ginderachter, Damya Laoui

**Author notes:** Equal contribution (Mohamed Lamkanfi, Jo A. Van Ginderachter, Damya Laoui). Corresponding authors: Damya Laoui, Jo A. Van Ginderachter.

## Abstract

Interleukin-1β (IL-1β) is a central mediator of inflammation whose secretion typically requires proteolytic maturation by the inflammasome and formation of membrane pores by gasdermin D (GSDMD). Emerging evidence suggests an important role for IL-1β in promoting cancer progression in patients, but the underlying mechanisms are little understood. Here, we show a key role for IL-1β in driving tumor progression in two distinct mouse tumor models. Notably, inflammasome activation and GSDMD were dispensable for the production of intratumoral bioactive IL-1β, which promoted systemic mobilization and infiltration of neutrophils into tumors. Neutrophils recruited via IL-1β suppressed the acquisition of an effector T-cell phenotype and subsequent antitumor immune response. Moreover, IL-1β was essential for neutrophil accumulation upon antiangiogenic therapy, thereby contributing to therapy-induced immunosuppression. Antitumor immunity in the absence of IL-1β-dependent neutrophil recruitment relied on immunostimulatory macrophages which promoted the infiltration and activation of cytotoxic T-cells. Overall, these results support a tumor-promoting role for IL-1β through establishing an immunosuppressive microenvironment and show that inflammasome activation is not essential for its release in tumors.

## INTRODUCTION

Chronic inflammation accompanying a developing tumor can promote its progression through various means, such as providing survival signals, suppressing T-cell function, inducing angiogenesis and enabling invasion and metastasis via tissue remodeling [1]. In addition, immune cells recruited to the tumor as part of the inflammatory response often antagonize anticancer therapies [2]. Hence, counteracting tumor-promoting inflammation appears to be key to improve disease outcome in many cancer types [3]. This, however, requires a more complete understanding of the mechanisms driving tumor-associated inflammation.

Interleukin-1β (IL-1β) is a pro-inflammatory cytokine whose role in cancer is increasingly recognized [4-6]. In a recent study, long-term treatment with a neutralizing anti-IL-1β antibody led to a dose-dependent reduction of lung cancer incidence and mortality in a large cohort of atherosclerosis patients with a history of myocardial infarction [7]. The presence of chronic IL-1β-driven inflammation in cancer was further supported by the identification of an IL-1β-induced transcriptional signature in the peripheral immune cells of renal cell cancer and breast cancer patients [8, 9]. Moreover, p53 mutations, late-stage disease and the basal-like subtype in breast cancer are all associated with significantly increased *IL1B* expression and may further augment systemic inflammation [9, 10]. IL-1β has been reported to promote tumor angiogenesis and the recruitment of myeloid cells, while it has also been shown to support antitumor T-cell responses and suppress metastatic outgrowth in mice, suggesting its role may be context-dependent [5, 6, 11-14]

IL-1β is produced as a biologically inactive precursor (pro-IL-1β) whose cleavage into the active form is typically mediated by caspase-1 [15]. Activation of caspase-1 is triggered by the inflammasome, a multiprotein complex which assembles upon activation of intracellular receptors, like NLRP3, AIM2 and NLRC4. Different inflammasome receptors are activated by distinct danger- or pathogen-associated molecular patterns, such as extracellular ATP, double-stranded DNA and bacterial flagellin [16]. Once cleaved, IL-1β follows an unconventional secretory pathway which typically requires membrane pores composed of gasdermin D (GSDMD) whose activation is also induced by the inflammasome [17].

Despite emerging interest in IL-1β as an oncology target, several questions regarding IL-1β signaling in the context of cancer remain unanswered. Firstly, although IL-1β secretion has been shown to be elevated in breast and lung tumors compared to adjacent non-involved tissues [9, 18], the exact cellular source of increased IL-1β production within these tumors remains poorly characterized. In fact, malignant cells, fibroblasts and immune cells have all been described as potential sources of IL-1β release in various tumor types [19-22].

Secondly, the inflammasome has been reported to be dispensable for IL-1β release in several types of sterile inflammation [23-25], raising the question whether caspase-1 is critically required for the proteolytic maturation and release of active IL-1β in the tumor microenvironment. GSDMD has been found to be essential for *in vivo* IL-1β release in several mouse models of inflammation, but it has not yet been determined whether it plays a similar role in tumors [26-29].

In addition, we still have a limited understanding about how IL-1β release impacts the complex tumor microenvironment and to what extent its effect is conserved across different tumor types. To address these knowledge gaps, we set out to examine the source of IL-1β in tumors, the requirement of inflammasomes and GSDMD for its release and its impact on the tumor microenvironment in mouse models of non-small cell lung cancer (NSCLC) and triple-negative breast cancer (TNBC).

## RESULTS

### Myeloid cells are the primary source of IL-1β in lung and breast tumors

In order to determine the cellular sources of IL-1β in lung and breast tumors with an unbiased approach, we analyzed single-cell RNA-seq datasets from human NSCLC and breast cancer. Unsupervised clustering of the data followed by identification of known cell lineages based on marker gene expression revealed 13 and 10 major cell types in lung and breast tumors, respectively (Figure 1A, Supplementary Figure 1). We found that the cell populations with the highest average expression levels of *IL1B* in both tumor types were myeloid cells, namely neutrophils, monocytes, dendritic cells (DCs) and macrophages, while other cell populations showed considerably (>10-fold) lower or no expression (Figure 1B,C). Of note, only a small number of neutrophils could be detected in these datasets, presumably due to their low transcript counts, while these cells are known to be well represented in both tumor types based on flow cytometry [30, 31].

**Figure 1.**
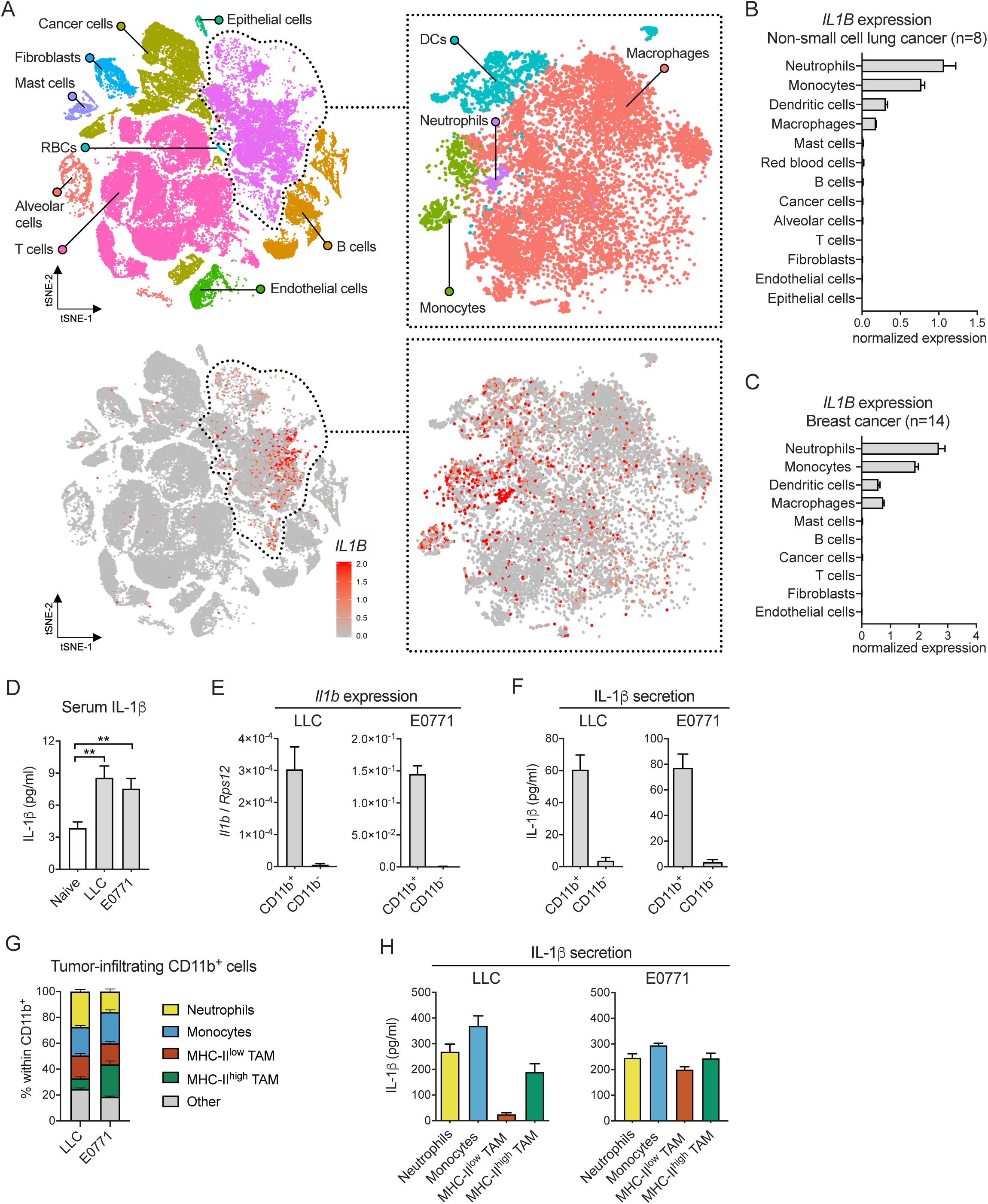
Myeloid cells are the primary source of IL-1β in lung and breast tumors. (A) tSNE plot of cells pooled from human lung tumors (n=8) color-coded based on cell clusters or *IL1B* expression level. (RBCs=Red blood cells, DCs=dendritic cells) (B) Average *IL1B* expression in different cell populations in human lung tumors based on pooled scRNA-seq data from 8 tumors. (C) Average *IL1B* expression in different cell populations in human breast tumors based on pooled scRNA-seq data from 14 tumors. (D) IL-1β concentration in the serum of naive mice (n=12) and mice with LLC (n=20) and E0771 (n=10) tumors. (E) Expression of *Il1b* measured by qPCR in the CD11b^+^ and CD11b^-^ fractions isolated by magnetic cell separation from LLC (n=4) and E0771 (n=7) tumors. (F) IL-1β secretion of the CD11b^+^ and CD11b^-^ fractions isolated by magnetic cell separation from LLC (n=3) and E0771 (n=5) tumors measured by ELISA after 24 h *in vitro* culture. (G) Relative frequency of major CD11b^+^ cell populations in LLC (n=10) and E0771 (n=11) tumors measured by flow cytometry. For gating strategies, see Supplementary Figure 2. (H) IL-1β secretion of the major CD11b^+^ cell populations isolated by cell sorting from LLC (n=4) and E0771 (n=3) tumors measured by ELISA after 24 h *in vitro* culture. Bar graphs show mean and SEM.

To investigate IL-1β production in more detail, we turned to mouse models of NSCLC and breast cancer, namely subcutaneous Lewis lung carcinoma (LLC), a p53-mutant lung adenocarcinoma, and orthotopic E0771, a p53-mutant TNBC model with basal-like characteristics [32-34]. Mice with LLC or E0771 tumors showed significantly elevated IL-1β levels in the serum compared to naive mice, indicating the presence of tumor-induced IL-1β-driven inflammation in these models (Figure 1D). To assess the contribution of myeloid cells to intratumoral IL-1β release, we separated the CD11b^+^ and CD11b^-^ fractions of tumors and measured the expression of *Il1b* mRNA in freshly isolated cells as well as the secretion of the cytokine following 24 h of *in vitro* culture. We found that both mRNA expression and protein secretion of IL-1β were almost exclusively restricted to the CD11b^+^ fraction of tumors (Figure 1E,F). Importantly, the LLC and E0771 cell lines did not show detectable IL-1β secretion *in vitro* (data not shown). The majority of the CD11b^+^ fraction in LLC and E0771 tumors consisted of neutrophils, monocytes and tumor-associated macrophages (TAMs) (Figure 1G, for gating strategies, see Supplementary Figure 2). Consistent with published reports, the TAM population included MHC-II^high^ and MHC-II^low^ subsets which possess immunostimulatory and anti-inflammatory gene expression profiles, respectively [35-38]. We then isolated these cell populations from tumors and assessed their IL-1β secretion *in vitro*. In LLC tumors, monocytes and neutrophils showed the highest secretion levels, followed by MHC-II^high^ TAMs and MHC-II^low^ TAMs (Figure 1H). In contrast, IL-1β secretion was comparable across the different myeloid cell types isolated from E0771 tumors (Figure 1H).

Altogether, these results demonstrate that myeloid cells are the primary source of IL-1β in human and mouse lung and breast tumors.

### IL-1β deletion inhibits tumor growth and reduces both systemic mobilization and tumor infiltration of neutrophils

To investigate the impact of IL-1β release on tumor progression, we implanted LLC or E0771 tumors in IL-1β-deficient (IL-1β^-/-^) mice and their wild-type (WT) littermates. Loss of IL-1β delayed tumor growth in both tumor types with a more pronounced effect in E0771 breast tumors where IL-1β-deficiency was, in some cases, associated with regression or durable tumor control (Figure 2A, Supplementary Figure 3A).

**Figure 2.**
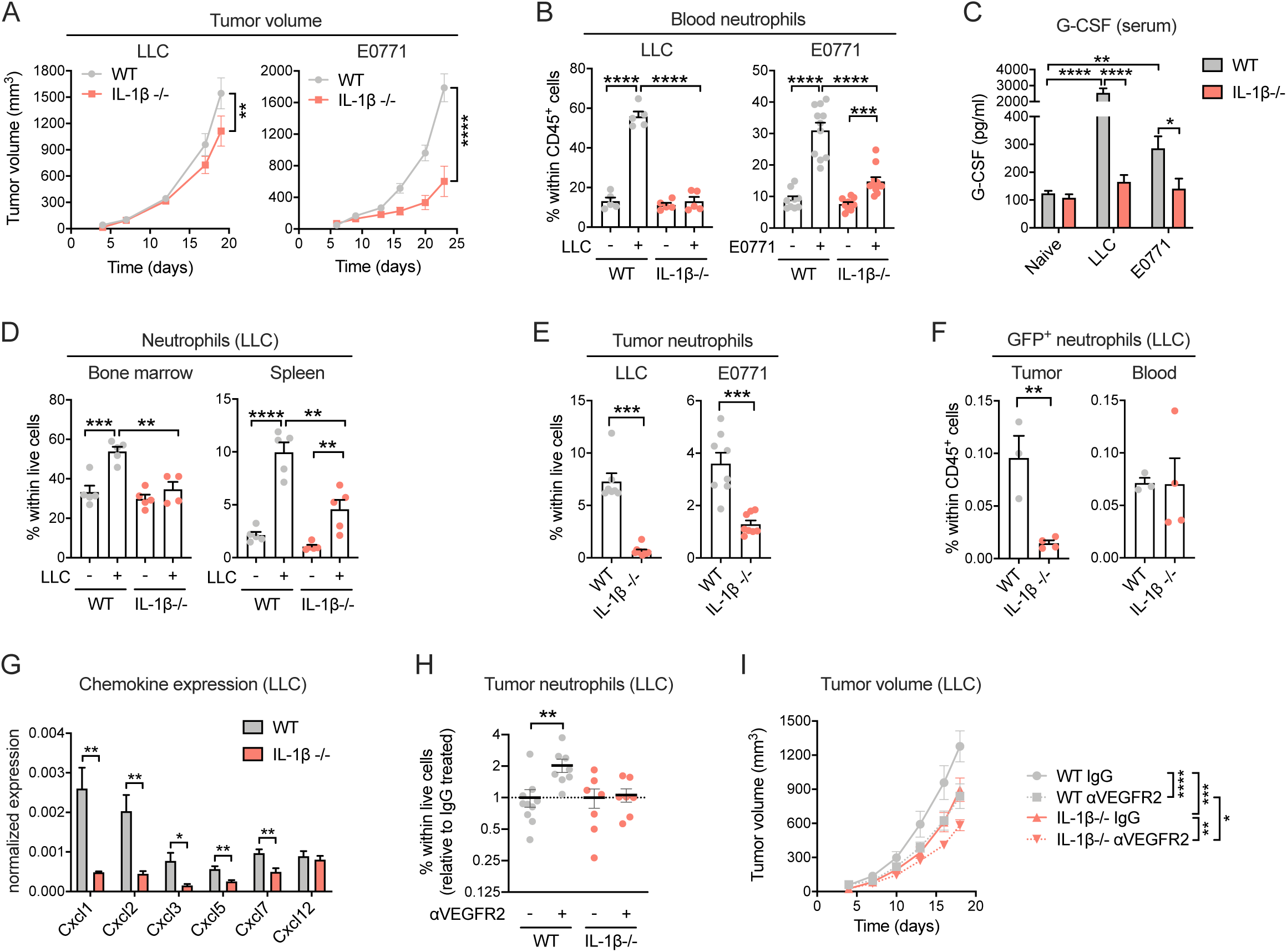
IL-1β deletion inhibits tumor growth and reduces both systemic mobilization and tumor infiltration of neutrophils. (A) LLC (n=9-10) and E0771 tumor growth (n=11) in WT and IL-1β-/- mice. (B) Frequency of neutrophils in the blood of naive and tumor-bearing WT and IL-1β-/- mice with LLC (n=5) and E0771 (n=10-11) tumors. (C) G-CSF levels measured via cytokine multiplex in the serum of naive (n=10) and tumor-bearing WT and IL-1β-/- mice with LLC (n=10) and E0771 (n=9-11) tumors. (D) Frequency of neutrophils in the bone marrow and spleen of naive and LLC tumor-bearing WT and IL-1β-/- mice (n=4-5). (E) Frequency of neutrophils in LLC (n=7-9) and E0771 (n=8) tumors in WT and IL-1β-/- mice. (F) Frequency of GFP^+^ neutrophils in LLC tumors and the blood 24 h after adoptive transfer in WT and IL-1β-/- mice (n=3-4). (G) Expression of neutrophil chemoattractants normalized to *Rps12* measured by qPCR in LLC tumor lysates of WT and IL-1β-/- mice (n=6-8). (H) Frequency of neutrophils measured by flow cytometry in LLC tumors of WT and IL-1β-/- mice treated with anti-VEGFR2 or IgG isotype control antibody normalized to the IgG isotype-treated control group within each genotype (n=7-10). (I) Volume of LLC tumors in WT and IL-1β-/- mice treated with anti-VEGFR2 or IgG isotype control antibody (n=8-10). Cell type frequencies were determined by flow cytometry. Graphs show mean and SEM.

IL-1β release induces neutrophilia during systemic inflammation and this may have an influence on tumor progression due to the wide range of tumor-promoting activities linked to neutrophils [39, 40]. For this reason, we analyzed the frequency of circulating CD11b^+^Ly6G^+^ neutrophils in naive and tumor-bearing WT and IL-1β^-/-^ mice. Both LLC and E0771 tumors induced expansion of circulating neutrophils which was abrogated in the absence of IL-1β (Figure 2B). These changes were mirrored by G-CSF levels in the blood, suggesting that tumor-induced, IL-1β-dependent neutrophil mobilization is driven by G-CSF (Figure 2C). Interestingly, loss of IL-1β prevented the LLC-induced expansion of bone marrow neutrophils, and partially also splenic neutrophils, suggesting that IL-1β-driven neutrophil expansion observed in the blood could be traced back to enhanced bone marrow output (Figure 2D). Next we assessed whether IL-1β-deficiency has an influence on neutrophils infiltrating primary tumors. We found that loss of IL-1β strongly reduced the abundance of neutrophils in both LLC and E0771 primary tumors (Figure 2E). To test whether the reduced abundance of tumor-infiltrating neutrophils is solely due to their decreased levels in the circulation or also due to altered recruitment, we adoptively transferred equal numbers of GFP-expressing splenic neutrophils into LLC tumor-bearing WT and IL-1β^-/-^ mice and assessed their frequency in the tumor after 24 hours. As shown in Figure 2F, recruitment of GFP^+^ neutrophils to the tumor was strongly reduced in IL-1β^-/-^ mice even though their frequency in the circulation was comparable to WT controls. Since CXCR2 ligands, particularly CXCL1 and CXCL2, have been shown to be critical for neutrophil extravasation [41], we asked whether these chemokines are affected by IL-1β release in the tumor. We found that all CXCR2 ligands, including CXCL1, CXCL2, CXCL3, CXCL5 and CXCL7, but not the CXCR4 ligand CXCL12, showed strongly reduced expression in the absence of IL-1β (Figure 2G).

Of note, the effect of IL-1β on the mobilization and recruitment of neutrophils was not restricted to the LLC and E0771 tumor models. We also observed a significant reduction of circulating and tumor-infiltrating neutrophils in IL-1β-deficient mice with EG7 lymphoma and B16-F10 melanoma tumors, which show greatly differing levels of neutrophil abundance (Supplementary Figure 3B,C).

Neutrophil recruitment to the tumor has been shown to drive therapy resistance and immunosuppression during treatment with antiangiogenic agents targeting VEGF signaling [42-45]. Hence, we wondered whether IL-1β is required for neutrophil infiltration during antiangiogenic therapy and examined the effect of anti-VEGFR2 antibody treatment in WT and IL-1β^-/-^ mice in the LLC tumor model. While anti-VEGFR2 treatment did not affect the levels of circulating neutrophils (Supplementary Figure 3D), we observed a 2-fold increase in the abundance of tumor-infiltrating neutrophils in treated WT mice, and this therapy-induced neutrophil recruitment was completely abrogated in IL-1β-deficient animals (Figure 2H). This was associated with a significantly reduced tumor burden in anti-VEGFR2-treated IL-1β^-/-^ mice compared to WT mice with the same treatment (Figure 2I).

Collectively, these results indicate that loss of IL-1β delays tumor progression in mouse models of NSCLC and TNBC and this is accompanied by reduced systemic mobilization and tumor infiltration of neutrophils. In addition, IL-1β-deletion prevents accumulation of neutrophils in the tumor triggered by anti-angiogenic therapy.

### Activation of the inflammasome and gasdermin D are dispensable for IL-1β-mediated neutrophil infiltration in tumors

Next, we asked whether the delayed tumor progression and strong reduction of neutrophil recruitment to tumors in IL-1β^-/-^ mice can be recapitulated in mice lacking various inflammasome components, which would suggest their requirement for bioactive IL-1β production in tumors. Deficiency of NLRP3 and NLRC4, two caspase-1-activating NOD-like receptors, did not affect *in vitro* IL-1β release of tumor-derived myeloid cells (Figure 3A; Supplementary Figure 4A). Accordingly, deletion of these inflammasome components did not alter tumor progression or neutrophil recruitment in mice with LLC and E0771 tumors as opposed to IL-1β deficiency (Figure 3B,C; Supplementary Figure 4B,C). In order to more directly assess the potential role of canonical and non-canonical inflammasome pathways, we next analyzed tumors in mice with combined deletion of inflammatory caspases 1 and 11 (Casp1/11^-/-^). Deletion of caspase-1/11 led to a partial reduction in IL-1β secretion levels by LLC tumor-derived myeloid cells (Figure 3D). However, this was not sufficient to alter tumor progression or neutrophil recruitment in LLC tumors (Figure 3E,F). IL-1β release by E0771 tumor-derived myeloid cells was not reduced in Casp1/11^-/-^ mice and, correspondingly, tumor growth and neutrophil infiltration remained unaltered in these tumors (Figure 3D-F). An IL-1β immunoblot on the culture supernatants of tumor-derived Casp1/11^-/-^ myeloid cells confirmed the inflammasome-independent production of mature IL-1β in both tumor models (Figure 3G). Caspase-8 has shown redundancy with caspase-1 in producing active IL-1β in some cases, cleaving pro-IL-1β at the same site [46, 47]. Active caspase-8 could be detected by immunoblot in sorted tumor-infiltrating myeloid cells but not in their circulating precursors (Supplementary Figure 4D). Hence, we assessed the contribution of caspase-8 to IL-1β release and neutrophil recruitment in tumors by using RIPK3^-/-^Casp8^-/-^ mice, in which RIPK3 deletion rescues embryonic lethality caused by caspase-8-deficiency [48]. We also generated Casp1/11^-/-^RIPK3^-/-^Casp8^-/-^ mice to evaluate the potential redundant roles of caspase-1/11 and -8. Caspase-8 deletion in both the RIPK3^-/-^ and Casp1/11^-/-^RIPK3^-/-^ backgrounds led to partial blockade of *in vitro* IL-1β release in myeloid cells derived from LLC tumors but not from E0771 tumors (Supplementary Figure 4E,H). However, this was not sufficient to affect tumor progression and neutrophil recruitment (Supplementary Figure 4F,G,I,J). Together, these data suggest slightly different mechanisms of IL-1β production by myeloid cells in LLC and E0771 tumors, but an overall independence of tumor growth and neutrophil recruitment from inflammasomes and caspase-8.

**Figure 3.**
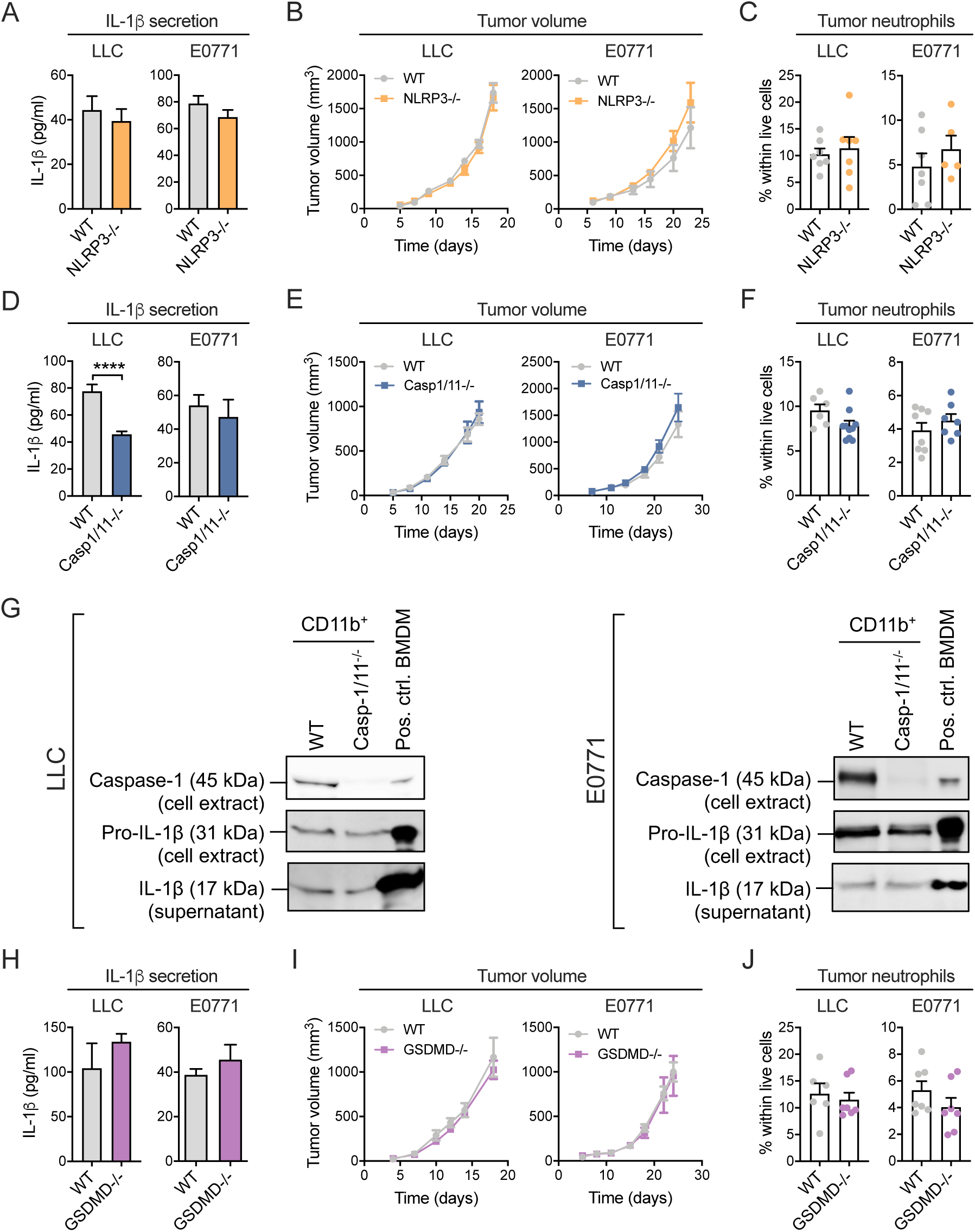
Activation of the inflammasome and gasdermin D are dispensable for IL-1β-mediated neutrophil infiltration in tumors. (A) IL-1β secretion of the CD11b^+^ fraction of LLC (n=7) and E0771 tumors (n=5-6) from WT and NLRP3-/- mice measured by ELISA following 24 h *in vitro* culture. (B) LLC (n=7) and E0771 (n=6-7) tumor growth in WT and NLRP3-/- mice. (C) Frequency of neutrophils in LLC (n=7) and E0771 (n=5-7) tumors in WT and NLRP3-/- mice. (D) IL-1β secretion of the CD11b^+^ fraction of LLC (n=7-8) and E0771 tumors (n=6-8) from WT and Casp1/11-/- mice measured by ELISA following 24 h *in vitro* culture. (E) LLC (n=5-6) and E0771 (n=8-9) tumor growth in WT and Casp1/11-/- mice. (F) Frequency of neutrophils in LLC (n=6-10) and E0771 (n=7-8) tumors in WT and Casp1/11-/- mice. (G) Immunoblots of caspase-1 and IL-1β on cell extracts and supernatants of CD11b^+^ myeloid cells isolated from LLC and E0771 tumors and cultured for 24 h *in vitro.* Mouse bone-marrow derived macrophages (BMDM) treated with LPS+ATP were used as positive controls. (H) IL-1β secretion of the CD11b^+^ fraction of LLC (n=4) and E0771 tumors (n=7-8) from WT and GSDMD-/- mice measured by ELISA following 24 h *in vitro* culture. (I) LLC (n=7) and E0771 (n=7) tumor growth in WT and GSDMD-/- mice. (J) Frequency of neutrophils in LLC (n=6-7) and E0771 (n=7) tumors in WT and GSDMD-/- mice. Neutrophil frequencies were determined by flow cytometry. Graphs show mean and SEM.

Membrane pore formation by GSDMD has proved critical for IL-1β release in mouse models of autoinflammation, steatohepatitis, disseminated intravascular coagulation and sepsis [26-29]. To investigate a potential role for GSDMD in IL-1β release, LLC and E0771 tumors were implanted in GSDMD^-/-^ mice and WT littermates. However, GSDMD-deficiency did not reduce IL-1β release of tumor-derived CD11b^+^ myeloid cells, tumor growth, and neutrophil recruitment (Figure 3H-J). Alternatively, necroptosis induced by membrane pores composed of mixed lineage kinase domain-like protein (MLKL) has been suggested to mediate IL-1β release independently of GSDMD-dependent pyroptosis *in vitro* [49]. However, neither MLKL-deficiency nor GSDMD/MLKL double-deficiency had a significant effect on myeloid cell IL-1β release, tumor progression and neutrophil recruitment (Supplementary Figure 5A-F).

Overall, these data from two distinct mouse tumor models demonstrate that activation of the inflammasome and caspase-8 as well as the formation of membrane pores by GSDMD and MLKL are dispensable for the release of bioactive IL-1β by tumor-associated myeloid cells and consequential neutrophil recruitment.

### IL-1β deletion enhances antitumor immunity

To investigate the mechanism of reduced tumor growth in IL-1β^-/-^ mice, we first turned to tumor angiogenesis, which has been described as being potentially IL-1β-regulated [11, 12]. However, we did not find any major differences in blood vessel density, pericyte coverage and vessel perfusion between IL-1β^-/-^ and WT mice (Supplementary Figure 6A-G). Slightly less hypoxic areas were observed in the tumors of IL-1β^-/-^ mice, however, this was likely due to the smaller average tumor size since volume-matched tumors did not show such difference (Supplementary Figure 6H). These observations suggested that IL-1β is not essential for tumor angiogenesis and therefore reduced tumor growth in IL-1β-deficient animals may be explained by alternative mechanisms.

Next, we examined whether neutrophils recruited by IL-1β to LLC or E0771 tumors were able to suppress T-cell proliferation. Indeed, CD11b^+^Ly6G^+^ neutrophils isolated from primary tumors were able to inhibit proliferation of splenocytes stimulated with anti-CD3 and anti-CD28 (Figure 4A). Tumor-infiltrating neutrophils from IL-1β^-/-^ mice showed similar T-cell suppressive activity to WT controls, suggesting that IL-1β dominantly affects their recruitment rather than their immunosuppressive activity (Figure 4B). In agreement with the immunosuppressive phenotype of tumor-infiltrating neutrophils, impaired neutrophil recruitment in IL-1β^-/-^ mice (Figure 2E) was accompanied by an elevated abundance of cytotoxic CD8^+^ T cells, while the infiltration of CD4^+^ T cells and FoxP3^+^ regulatory T cells remained unaltered (Figure 4C-E). Furthermore, a higher proportion of tumor-infiltrating CD4^+^ and CD8^+^ T cells showed an effector T-cell phenotype in IL-1β^-/-^ mice as indicated by the increased CD44^+^CD62L^-^ effector vs. CD44^-^CD62L^+^ naive T-cell ratio (Figure 4F,G, Supplementary Figure 7A). Correspondingly, enhanced infiltration of neutrophils upon anti-VEGFR2 therapy in WT mice (Figure 2H) was associated with impaired effector T-cell differentiation and this was counteracted by IL-1β deletion (Figure 4H, Supplementary Figure 7B). Among the immune cells possessing T-cell stimulatory potential, conventional DCs did not show altered infiltration in tumors of IL-1β^-/-^ mice (Supplementary Figure 7C). Although we observed decreased infiltration of monocytes in LLC tumors of IL-1β^-/-^ hosts (Figure 4I), TAM abundance was not reduced in neither LLC nor E0771 tumors (Figure 4J). However, we found higher abundance of MHC-II^high^ TAMs (Figure 4K), a TAM phenotype which has been shown to be driven by effector T cells [50-52], and, in turn, possesses the capacity to stimulate T-cell responses [35, 38, 51]. In line with these observations, presence of MHC-II^high^ TAMs but not MHC-II^low^ TAMs showed a strong positive correlation with effector T-cell infiltration in both LLC and E0771 tumors (Figure 4L, Supplementary Figure 7D), suggesting that the expansion of MHC-II^high^ TAMs may participate in amplifying the antitumor T-cell response in the absence of IL-1β.

**Figure 4.**
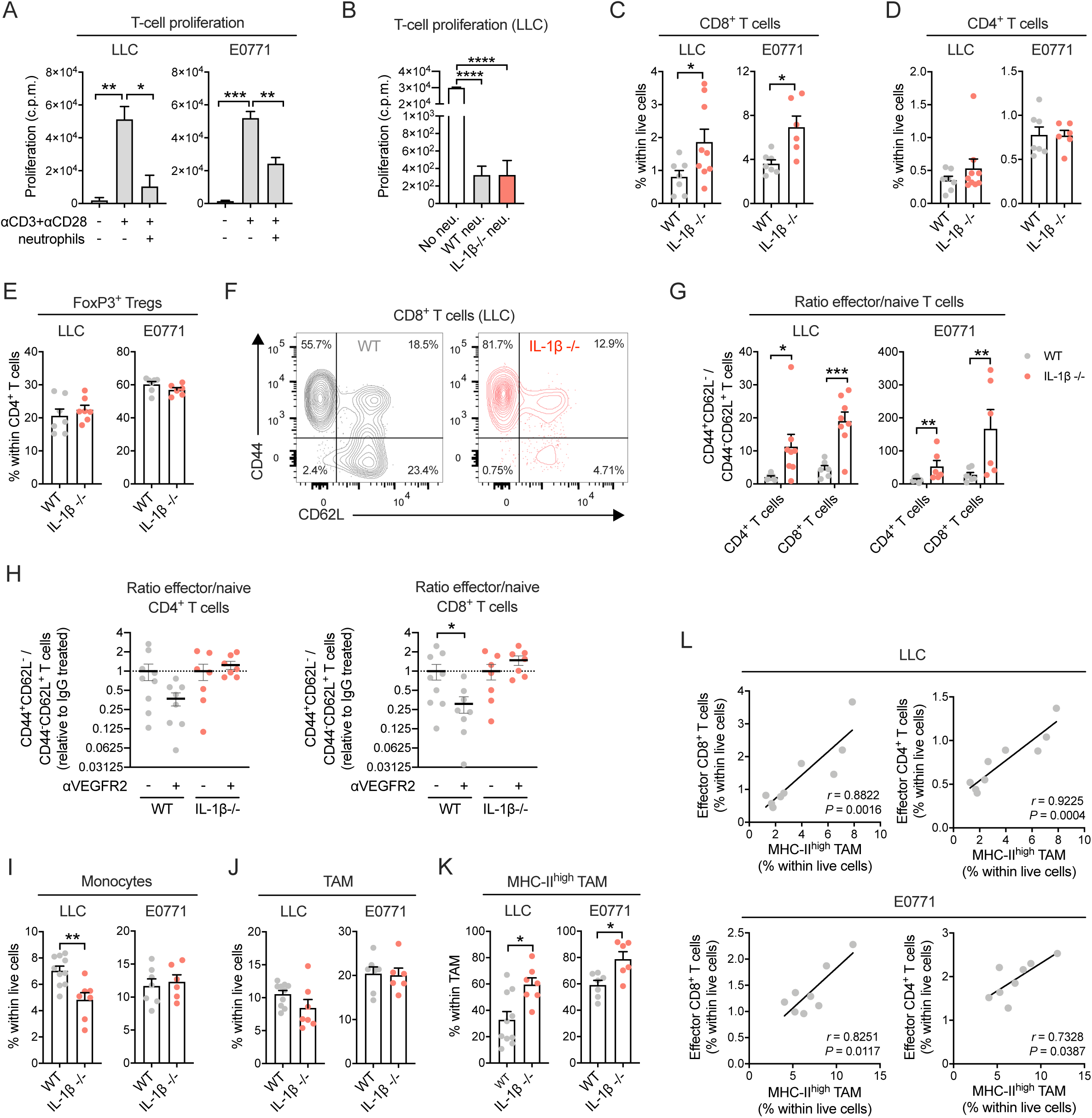
IL-1β deletion enhances antitumor immunity. (A) T cell proliferation following co-culture of splenocytes with LLC/E0771 tumor-derived neutrophils in a 1:1 ratio, measured via ^3^H-thymidine incorporation (c.p.m.: count per minute). (n=3, data pooled from 3 independent experiments). (B) T cell proliferation following co-culture of splenocytes with LLC tumor-derived WT and IL-1β -/- neutrophils in a 1:1 ratio, measured via ^3^H-thymidine incorporation (c.p.m.: count per minute). (n=3, data pooled from 3 independent experiments). (C)-(E) Frequency of the indicated cell populations in LLC (n=7-9) and E0771 (n=6-7) tumors of WT and IL-1β -/- mice. (F) Representative flow cytometry plot of CD44 and CD62L expression on CD8^+^ T cells in LLC tumors of WT and IL-1β -/- mice. (G) Ratio of effector (CD44^+^CD62L^-^) and naive (CD44^-^CD62L^+^) T cells in LLC (n=6-8) and E0771 (n=6-7) tumors of WT and IL-1β -/- mice. For frequencies of effector and naive T cells see Supplementary Figure 6A. (H) Ratio of effector (CD44^+^CD62L^-^) and naive (CD44^-^CD62L^+^) T cells measured by flow cytometry in LLC tumors of WT and IL-1β-/- mice treated with anti-VEGFR2 or IgG isotype control antibody normalized to the IgG isotype-treated control group within each genotype (n=7-9). For frequencies of naive and effector T cells, see Supplementary Figure 7B. (I)-(K) Frequency of the indicated cell populations in LLC (n=7-10) and E0771 (n=6-7) tumors of WT and IL-1β -/- mice. (L) Correlation of CD44^+^CD62L^-^ effector T cell and MHC-II^high^ TAM abundance in LLC (n=9) and E0771 (n=8) tumors. Pearson *r* values and *P* values are indicated in the graphs. Cell type frequencies were determined by flow cytometry. For gating strategies see Supplementary Figure 2. Bar graphs show mean and SEM.

In summary, these data indicate a crucial role for IL-1β in promoting an immunosuppressive tumor microenvironment driven by the recruitment of neutrophils, leading to impaired accumulation of effector T cells and inhibition of antitumor immunity.

### Antitumor immunity in IL-1β-deficient mice is macrophage-dependent

In order to test the potential contribution of TAMs to the enhanced antitumor cytotoxic T-cell response observed in IL-1β-deficiency, we set out to deplete these cells in tumor-bearing mice using the CSF1R inhibitor PLX5622. This small-molecule inhibitor is highly specific for CSF1R and has been successfully used before to deplete TAMs [53]. Administration of PLX5622 reduced TAM infiltration in LLC tumors by 89%, whereas it only caused a 28% reduction in E0771 tumors (Supplementary Figure 8A). Based on these results, we decided to examine the impact of TAM depletion in the LLC tumor model. Depletion of TAMs in IL-1β^-/-^ mice restored tumor growth to WT levels, while it did not have an effect in WT mice (Figure 5A). Analysis of the tumor immune cell composition confirmed that CSF1R inhibition efficiently eliminated TAMs in both WT and IL-1β^-/-^ mice while it did not deplete neutrophils, monocytes and DCs in tumors (Figure 5B,C, Supplementary Figure 8B). Analysis of tumor-infiltrating T-cells revealed that TAM depletion in IL-1β^-/-^ mice reduced CD8^+^ T-cell abundance to the WT level (Figure 5D). Similarly, TAM depletion in IL-1β^-/-^ mice restored the ratio of tumor-infiltrating effector vs. naive CD8^+^ T cells to levels similar to WT (Figure 5E, Supplementary Figure 8C). In addition, the proportions of CD8^+^ T cells expressing the activation markers CD69 and granzyme B were reduced by TAM depletion in both WT and IL-1β^-/-^ mice (Figure 5F-G). In accordance with these results, TAM depletion led to lower intratumoral expression of chemokines commonly associated with T-cell trafficking, including CXCL9, CXCL10, CXCL16 and CCL5 (Figure 5H) [54]. TAM depletion in IL-1β^-/-^ mice also reduced the intratumoral expression of the costimulatory molecules CD40 and CD86 and the Th1 stimulatory cytokine IL-12, further supporting that TAMs in LLC tumors are an important source of T-cell stimulatory signals (Figure 5I).

**Figure 5.**
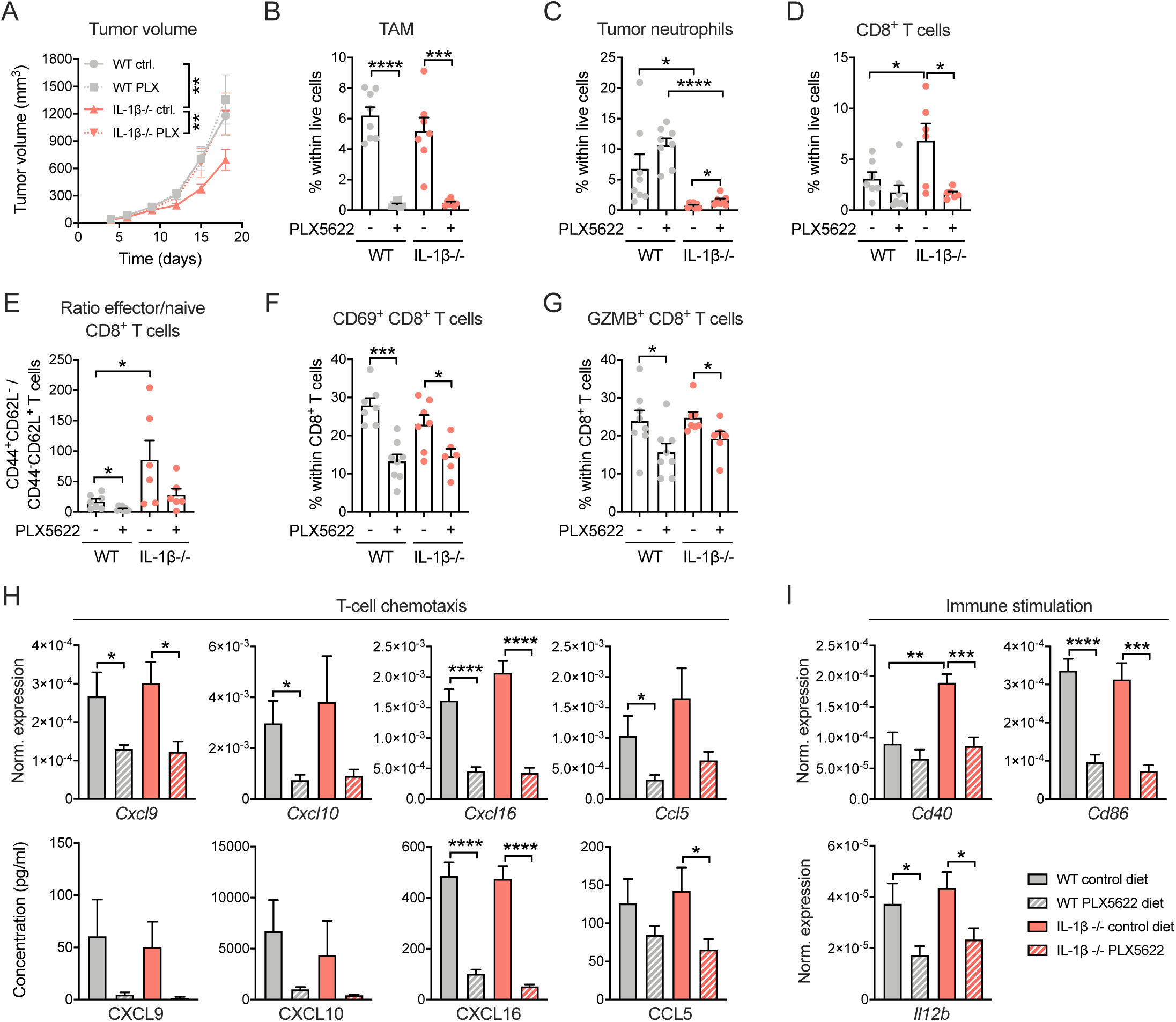
Antitumor immunity in IL-1β-deficient mice is macrophage-dependent. (A) LLC tumor growth in WT and IL-1β-/- mice treated with PLX5622 or control diet (n=7-8). (B)-(G) Frequency of indicated cell populations in LLC tumors of WT and IL-1β-/- mice treated with PLX5622 or control diet measured by flow cytometry (n=7-8). (H) Expression of selected chemokines normalized to *Rps12* measured by qPCR in LLC tumor lysates of WT and IL-1β-/- mice treated with PLX5622 or control diet (n=7-8) and concentration of the same chemokines measured by ELISA/multiplex immunoassay in total tumor supernatants after 24 h *in vitro* culture. (I) Expression of selected immunostimulatory genes normalized to *Rps12* measured by qPCR in LLC tumor lysates of WT and IL-1β-/- mice treated with PLX5622 or control diet (n=7-8). Graphs show mean and SEM.

Together, these results suggest that TAMs support the accumulation and activation of CD8^+^ T cells in tumors in the absence of IL-1β, thereby contributing to an antitumor immune response.

## DISCUSSION

In this study, we demonstrate in two distinct mouse models that IL-1β, released mainly by neutrophils, monocytes and macrophages, plays a key role in mobilizing neutrophils from the bone marrow during tumor progression and induces their infiltration into tumors. Our results are in line with previous reports demonstrating the role of IL-1β in systemic neutrophil mobilization in breast cancer, however, these studies only focused on its consequences on the metastatic environment and not the primary tumor [14, 20]. Earlier studies utilizing IL-1β blockade or IL-1β-overexpressing cancer cells have demonstrated that this cytokine promotes infiltration of myeloid cells into tumors, but the exact identity of these cells remained unknown [11, 55, 56]. More recently, IL-1β was reported to induce CCL2 and promote the recruitment of monocytes and subsequent accumulation of macrophages in the 4T1 mouse model of breast carcinoma [57]. However, we did not observe a similar impairment of monocyte recruitment in the absence of IL-1β in orthotopic E0771 breast tumors, suggesting that the link between IL-1β release and monocyte recruitment is not a general phenomenon.

High neutrophil-to-lymphocyte ratio correlates with worse overall survival in both lung and breast cancer, suggesting that neutrophil expansion associated with a systemic inflammatory response may be an important determinant of disease outcome [58, 59]. In addition, intratumoral neutrophil abundance inferred from bulk tumor transcriptome data shows the strongest negative correlation with patient prognosis in lung and breast tumors across 25 different tumor types [60]. A growing body of evidence from mouse studies adds support to a causal link between neutrophils and tumor progression in various tumor types [40]. Two major ways through which neutrophils can facilitate tumor growth are the induction of angiogenesis and suppression of antitumor immunity [40]. In the current study, we did not observe any quantitative or qualitative changes in the vasculature of LLC tumors implanted in IL-1β-deficient mice in which neutrophils were virtually absent from tumors. It is conceivable that neutrophils do not possess proangiogenic activity in the tumor types examined here or their loss is compensated by the strongly proangiogenic macrophages present [36]. In contrast, we observed evidence for enhanced antitumor immunity in the absence of IL-1β-driven immunosuppressive neutrophil recruitment in both LLC and E0771 tumors which was associated with reduced primary tumor growth and even tumor regression in some animals in the case of E0771 tumors. These results are in agreement with previous studies which showed that neutrophil depletion by anti-Ly6G antibody or CXCR2 blockade led to increased abundance of effector T cells in tumors [20, 61]. The difference in the extent of tumor growth inhibition between the two models may be explained by the fact that LLC is considered poorly immunogenic, while E0771 has a high mutation and predicted neoantigen burden and responds well to immunotherapy [34, 62].

A notable observation in this study is that macrophage depletion led to the reversal of the enhanced antitumor immunity in IL-1β^-/-^ mice, suggesting the presence of immunostimulatory TAMs in tumors. Although TAMs are believed to primarily act as tumor-supporting cells, this cell type shows considerable heterogeneity in tumors and the presence of immunostimulatory macrophage populations has been demonstrated in both mouse and human tumors [63-65]. In agreement with our findings, previous studies have shown that the immunostimulatory TAM population expands upon T-cell activation and amplifies cytotoxic T-cell responses [51, 66, 67].

We also show in this study that the canonical and non-canonical inflammasomes are dispensable for the production of bioactive IL-1β in LLC and E0771 tumors, and combined deletion of caspase-1/11 was not sufficient to recapitulate the *in vivo* phenotype observed in IL-1β^-/-^ mice. Although caspase-8 was activated in tumor-infiltrating myeloid cells, deletion of this enzyme was not sufficient to completely block bioactive IL-1β release and neutrophil infiltration. A diverse range of additional enzymes have been shown to cleave pro-IL-1β, including proteinase 3, neutrophil elastase, cathepsin G, granzyme A, chymase, matrix metalloproteinases and meprins [15, 68, 69]. Several of the aforementioned enzymes may be active and play redundant roles in the tumor microenvironment, therefore it might not be possible to pinpoint a single enzyme which is responsible for IL-1β production in tumors. Since several reports have demonstrated the beneficial effect of genetic or pharmacological inhibition of NLRP3 and caspase-1 on tumor progression in mice, it is likely that requirement of the inflammasome for IL-1β release depends on the availability of alternative cleavage pathways determined by the immune microenvironment [6]. Alternatively, decreased tumor progression observed in some of these studies may also be explained by IL-1β-independent effects such as inhibition of inflammasome-mediated IL-18 release [16].

To our knowledge, the contribution of GSDMD and MLKL to IL-1β release in tumors had not been evaluated before. In LLC and E0771 tumors, these pore-forming proteins were not required for IL-1β release and neutrophil recruitment. This suggests the existence of alternative release mechanisms that may include passive release through myeloid cell necrosis which has been linked to IL-1β release *in vitro* and is likely to occur in the tumor microenvironment [70]. Altogether, our data suggest a model whereby IL-1β released from tumor-infiltrating myeloid cells increases serum G-CSF levels and intratumoral expression of CXCR2 ligands, promoting systemic neutrophil mobilization and extravasation in the tumor, respectively. Recruited neutrophils exhibit immunosuppressive activity and impede the acquisition of an effector T-cell phenotype. This, in turn, dampens activation of TAMs which are required for the recruitment and activation of T cells. Administration of VEGFR2-blocking antiangiogenic therapy exacerbates this process, further increasing neutrophil infiltration and consequently reducing effector T-cell accumulation in an IL-1β-dependent manner. Collectively, these observations provide support for the role of IL-1β as a tumor-promoting factor whose inactivation results in an immune permissive tumor microenvironment. We suggest that the existence of inflammasome-independent IL-1β release and neutrophil recruitment demonstrated here will have to be taken into consideration when applying the growing range of inflammasome inhibitors for cancer therapy [6].

## MATERIALS AND METHODS

### Mice

All experiments were performed with age-matched female mice. C57BL/6 mice were from Janvier, IL-1β^-/-^ mice were provided by François Huaux (UCL, Belgium), UBI-GFP mice were from Jackson. NLRP3^-/-^ [71], NLRC4^-/-^ [72], Casp1/11^-/-^ [73], RIPK3^-/-^Casp8^-/-^ [48], GSDMD^-/-^ [29] and MLKL^-/-^ [74] mice were described previously. Casp1/11^-/-^RIPK3^-/-^Casp8^-/-^ and GSDMD^-/-^MLKL^-/-^ mice were generated in the VIB Center for Inflammation Research, Ghent, Belgium. In all experiments involving knock-out mice, WT (+/+) or heterozygote (+/-) littermate mice were used as controls as specified in the figures and figure legends.

All procedures followed the guidelines of the Belgian Council for Laboratory Animal Science and were approved by the Ethical Committee for Animal Experiments of the Vrije Universiteit Brussel (licenses 14-220-26, 16-220-3, 19-220-35).

### Tumor models

LLC, EG7 and B16F10 cell lines were from ATCC, E0771 cells were from CH3Biosystems. LLC, E0771 and B16F10 cells were maintained in DMEM supplemented with 10% (v/v) heat-inactivated fetal calf serum (FCS; Capricorn Scientific), 300 μg/ml L-glutamine, 100 units/ml penicillin and 100 μg/ml streptomycin. For EG7 cells, DMEM was replaced by RPMI.

For tumor implantation, 3×10^6^ LLC cells, 1×10^6^ B16F10 cells or 3×10^6^ EG7 cells were injected subcutaneously into the right flank of mice in 200 μl of HBSS. For orthotopic breast tumor implantation, 5×10^5^ E0771 cells were injected into the left 4th mammary fat pad in 50 μl of HBSS mixed with Growth Factor Reduced Matrigel (Corning) in a 1:1 ratio.

Tumor volumes were determined by caliper measurements and calculated using the formula: V = π × (d^2^ × D)/6, where d is the shortest diameter and D is the longest diameter.

### Treatments

CSF1R inhibitor PLX5622 was administered via rodent chow (1200 mg PLX5622/kg chow) starting from day 6 of tumor growth. PLX5622 was provided by Plexxikon. Control and PLX5622-containing AIN-76A rodent chow was prepared by Research Diets.

Anti-VEGFR2 (clone DC101, BioXCell) or isotype control antibody (clone HRPN, BioXCell) was administered intraperitoneally every 3 days starting from day 4 of tumor growth at a dose of 40 mg/kg body weight.

### Blood collection and tissue dissociation

Blood was collected in tubes containing EDTA. Tumors were excised, cut in small pieces, incubated with 10 U/ml collagenase I, 400 U/ml collagenase IV and 30 U/ml DNase I (Worthington) in RPMI for 30 min at 37 °, squashed and filtered. Spleens were mashed through a cell strainer, bone marrow was flushed out from the femurs into RPMI. All single-cell suspensions were treated with ACK (Ammonium-Chloride-Potassium) erythrocyte lysis buffer.

### Flow cytometry and cell sorting

Single cell suspensions were resuspended in HBSS and samples for flow cytometry analysis were incubated with Fixable Viability Dye eFluor 506 (1:1000, eBioscience) for 30 min at 4 °C. Next, cell suspensions were washed with HBSS and resuspended in HBSS with 2 mM EDTA and 1% (v/v) FCS. To prevent nonspecific antibody binding to Fcγ receptors, cells were pre-incubated with anti-CD16/CD32 (clone 2.4G2) antibody. Cell suspensions were then incubated with fluorescently labelled antibodies diluted in HBSS with 2 mM EDTA and 1% (v/v) FCS for 20 min at 4°C and then washed with the same buffer. The following fluorochrome-conjugated antibody clones were used: CD45 (30-F11), CD11b (M1/70), Ly6G (1A8), SiglecF (E50-2440), MHC-II (M5/114.15.2), Ly6C (HK1.4), F4/80 (CI:A3-1), CD11c (HL3), CD24 (M1/69), NK1.1 (PK136), CD19 (1D3), TCRβ (H57-597), CD4 (RM4-5), CD8 (53-6.7), FoxP3 (FJK-16s), CD44 (IM7), CD62L (MEL-14), CD69 (H1.2F3), GZMB (GB11), CD31 (390).

Flow cytometry data were acquired using a BD FACSCanto II (BD Biosciences) and analyzed using FlowJo. The gating strategy to identify immune cell populations in tumors is shown in Supplementary Figure 2. Samples with less than 10% viable cells and tumor samples with cell contamination from the tumor-draining lymph node (identified as outliers in B cell and naive T cell abundance) were excluded from further analyses.

For fluorescence-activated cell sorting of myeloid cell populations, tumor single cell suspensions were enriched for CD11b^+^ cells using magnetic cell separation (Miltenyi). 7-AAD staining was used to exclude dead cells. Cells subsets were then sorted into RPMI with 10% (v/v) FCS, 300 μg/ml L-glutamine, 100 units/ml penicillin, 100 μg/ml streptomycin, 1% (v/v) MEM non-essential amino acids (11140050, Gibco), 1 mM sodium pyruvate and 0.02 mM 2-mercaptoethanol. Fluorescence-activated cell sorting was performed using a BD FACSAria II (BD Biosciences).

Magnetic cell separation of CD11b^+^ myeloid cells was performed according to the manufacturer’s protocol (Miltenyi).

### *Ex vivo* cell culture

The viability of cells after cell sorting was confirmed using trypan blue staining. For *ex vivo* culture, 3×10^5^ cells/well were cultured for 24 h in flat-bottom 96 well plates in 200 μl/well RPMI containing 10% (v/v) FCS, 300 μg/ml L-glutamine, 100 units/ml penicillin, 100 μg/ml streptomycin, 1% (v/v) MEM non-essential amino acids (11140050, Gibco), 1 mM sodium pyruvate and 0.02 mM 2-mercaptoethanol.

### Adoptive transfer of neutrophils

Neutrophils from the spleen of LLC tumor-bearing UBI-GFP mice were isolated by magnetic cell separation using anti-Ly6G microbeads according to the manufacturer’s protocol (Miltenyi). 5×10^6^ GFP-expressing neutrophils in 100 μl HBSS were injected through the tail vein into recipient LLC tumor-bearing mice which were sacrificed 24 h later.

### T-cell suppression assay

2×10^5^ neutrophils sorted from tumors were added to 2×10^5^ naïve C57BL/6 splenocytes stimulated with anti-CD3 (1 μg/ml) and anti-CD28 (2 μg/ml) and cultured in flat-bottom 96-well plates in culture medium for ex vivo cell culture described above. After 24 h of culture, 1 μCi (0.037 MBq) ^3^H-thymidine was added and after another 18 h of culture T-cell proliferation was measured as count per minute in a liquid scintillation counter.

### RNA extraction, cDNA preparation, and quantitative real-time PCR

Tumor tissue was snap-frozen in liquid nitrogen and homogenized in 1 ml TRIzol (Invitrogen) in gentleMACS M tubes using the gentleMACS Dissociator (Miltenyi). RNA was extracted using TRIzol (Invitrogen) and was reverse-transcribed with oligo(dT) and SuperScript II RT (Invitrogen) following the manufacturers’ protocols. Quantitative real-time PCR was performed in the CFX Connect Real-Time System (Bio-Rad) using the SsoAdvanced Universal SYBR Green Supermix (Bio-Rad) and the following primers: *Rps12*-F: GGAAGGCATAGCTGCTGGAGGTGT, *Rps12*-R: CCTCGATGACATCCTTGGCCTGAG; *Il1b*-F: GTGTGGATCCCAAGCAATAC, *Il1b*-R: GTCTGCTCATTCACGAAAAG; Cxcl1-F: GCTTGAAGGTGTTGCCCTCAG, Cxcl1-R: AAGCCTCGCGACCATTCTTG; Cxcl2-F: TGGAAGGAGTGTGCATGTTC, Cxcl2-R: CACGAAAAGGCATGACAAAA; Cxcl3-F: CACCCAGACAGAAGTCATAGCCAC, Cxcl3-R: TGGTGAGGGGCTTCCTCCTTT; Cxcl5-F: CTCGCCATTCATGCGGAT, Cxcl5-R: CTTCAGCTAGATGCTGCGGC; Cxcl7-F: CTCAGACCTACATCGTCCTGC, Cxcl7-R: GTGGCTATCACTTCCACATCAG; Cxcl12-F: TCATCCCCATTCTCCTCATC, Cxcl12-R: ATAAAGGAGCCTCCCTCTGC; *Cxcl9*-F: CCTCCTTGCTTGCTTACCAC, *Cxcl9*-R: TTTTCACCCTGTCTGGCTCT; *Cxcl10*-F: AATTGCCCTTGGTCTTCTGA, *Cxcl10*-R: CCTTGGGAAGATGGTGGTTA; *Cxcl16*-F: GTCTCCTGCCTCCACTTTCT, *Cxcl16*-R: CTAAGGGCAGAGGGGCTATT; *Ccl5*-F: GTGCCCACGTCAAGGAGTAT, *Ccl5*-R: CGAGTGGGAGTAGGGGATTA; *Il12b*-F: TCAGGGACATCATCAAACCA, *Il12b*-R: CTACGAGGAACGCACCTTTC; *Cd40*-F: GCTGTGAGGATAAGAACTTGGAGG, *Cd40*-R: GCATCCGGGACTTTAAACCACA; *Cd86*-F: CCTCCAAACCTCTCAATTTCA, *Cd86*-R: TCGGCTTCTTGTGACATACAAT. The following program was used for real-time PCR: 95 °C 3 min, 40×(94 °C 30 s, 54 °C 30 s, 72 °C 45 s). Gene expression was normalized to *Rps12*.

### Cytokine measurements

IL-1β and CXCL9 were measured using ELISA (IL-1β from cell culture supernatants: MLB00C, R&D Systems; IL-1β from serum: MHSLB00, R&D Systems; CXCL9: DY492, R&D Systems), G-CSF, CXCL10, CXCL16 and CCL5 were measured using multiplex immunoassay (Bio-Rad) according to the manufacturers’ protocols.

### Western blotting

Cell lysates for immunoblots were prepared by resuspending cells in a lysis buffer containing 20 mM Tris HCl (pH 7.4), 200 mM NaCl and 1% (v/v) NP-40. Cell lysates and cell culture supernatants were denatured in Laemmli buffer at 95°C for 10 min. SDS-PAGE–separated proteins were transferred to PVDF membranes. Blocking, incubation with antibody, and washing of the membrane were done in PBS supplemented with 0.05% (v/v) Tween 20 and 3% (v/v) non-fat dry milk. Immunoblots were incubated overnight with primary antibodies against caspase-1 (AG-20B-0042-C100, Adipogen) and IL-1β (GTX74034, Genetex). Horseradish peroxidase-conjugated goat anti-mouse (115-035-146, Jackson ImmunoResearch Laboratories) or anti-rabbit (111-035-144, Jackson ImmunoResearch Laboratories) secondary antibody was used to detect proteins by enhanced chemiluminescence (Thermo Scientific). Mouse bone marrow-derived macrophages (BMDMs) treated with 0.5 μg/ml LPS (tlrl-smlps, Invivogen) for 3 h followed by 5 mM ATP (10519987001, Roche) for 45 min were used as positive controls for caspase-1 and IL-1β blots. BMDMs were generated by culturing bone marrow cells in IMDM (Lonza) containing 10% (v/v) FCS, 30% (v/v) L929 cell-conditioned medium, 1% (v/v) MEM non-essential amino acids (Lonza), 100 U/ml penicillin, and 100 μg/ml streptomycin at 37°C in a humidified incubator containing 5% CO_2_ for 6 days.

### Histology

For the assessment of tumor blood vessel perfusion, mice were injected intravenously with 0.05 mg FITC-conjugated lectin (Lycopersicon esculentum; Vector Laboratories). After 10 minutes, mice were sacrificed and tumors were harvested.

Tumor hypoxia was detected via intraperitoneal injection of 60 mg/kg body weight pimonidazole hydrochloride (Hypoxyprobe) into tumor-bearing mice. After 1 h, mice were sacrificed and tumors were harvested.

Tumor samples were fixed in 2% PFA overnight at 4°C, then dehydrated and embedded in paraffin. Serial sections of 7 μm thickness were made. Slides were first rehydrated to further proceed with antigen retrieval in citrate solution (DAKO) at 100 °C for 20 min. Slides were then incubated in 0.3% hydrogen peroxide in methanol for 20 min to block endogenous peroxidases. The sections were blocked with donkey serum (Sigma) for 45 min and incubated overnight at room temperature with the following antibodies: anti-CD31 (550274, BD Biosciences), anti-FITC (4510-7604, Serotec), anti-αSMA-Cy3 (C6198, Sigma), anti-pimonidazole (4.3.11.3, Hypoxyprobe). Next, appropriate secondary Alexa Fluor 488/647-conjugated antibodies (Invitrogen) or biotin-labeled antibodies (Jackson Immunoresearch) were applied. After biotin-labelled antibodies, TSA Cyanine 3 or Cyanine 5 amplification kits (Perkin Elmer) were used according to the manufacturer’s instructions. Hoechst solution was used to stain nuclei. Mounting of slides was done with ProLong Gold mounting medium without DAPI (Invitrogen). Imaging and microscopic analysis was performed with an Olympus BX41 microscope and CellSense imaging software. Slides were scanned using Zeiss AxioScan Z.1 slide scanner. CD31^+^ blood vessel density and the proportion of FITC-lectin^+^ (perfused) and αSMA^+^ (pericyte-covered) blood vessels were determined by manual counting in 6 representative microscopic images/tumor. The proportion of pimonidazole^+^ hypoxic areas were determined in whole tumor cross-sections using ImageJ.

### Analysis of single-cell RNA-seq data from human tumors

The droplet-based scRNA-seq data of 8 untreated lung cancer patients [75] (10x Genomics 3’ RNA library kit, ArrayExpress:E-MTAB-6149 and E-MTAB-6653) were processed and clustered using Seurat (v2.3.4) package. Cell matrix was filtered (nUMI > 400, 200 < nGene < 6000, mitochondrial RNA < 25%), normalized, regressed for confounding factors (nUMI, patient, mitochondrial RNA and cell cycle) and scaled. The variable genes (normalized expression between 0.125 and 3, quantile-normalized variance > 0.5) were used to construct principal components (PCs), followed by graph-based clustering (tSNE and Louvain algorithm). Cell type annotation was based on the expression of established marker genes. pDCs were initially co-clustered with B-cells, and then annotated back to myeloid population, where most other DCs were co-clustered with. Then the myeloid cells were subclustered to identify monocytes (*SELL, CDKN1C, MTSS1*), macrophages (*CD68, CD163, MCR1*), DCs (*CLEC9A, XCR1, CD1C*, CD1A, *LILRA4*) and neutrophils (*FCGR3B*). Similar analysis was performed for 5’-scRNA-seq data from 14 treatment-naïve breast cancers [76], and the myeloid cells were further subclustered and annotated.

### Statistical analysis

Statistical analyses were performed in GraphPad Prism software. For relevant pairwise comparisons, unpaired two-tailed t-test was used to calculate the *P* value. Tumor growth curves were compared by 2-way ANOVA with Holm-Sidak multiple comparisons test. To assess correlation, Pearson correlation coefficient was calculated. A *P* value < 0.05 was considered statistically significant. For statistically significant differences, the *P* value is indicated in graphs as the following: * *P*<0.05, ** *P*<0.01, *** *P*<0.001, **** *P* < 0.0001. Comparisons found to be nonsignificant are not shown.

## ACKNOWLEDGEMENTS

We thank Dr. Geert Van Loo, Dr. Lars Vereecke and Mozes Sze for help with the multiplex immunoassays. We thank Maryse Schmoetten, Jan Brughmans, Lea Brys, Ella Omasta, Marie-Therese Detobel and Nadia Abou for technical and administrative assistance. We would like to thank the VIB BioImaging Core for training, support and access to the instrument park and Amanda Gonçalves for help with slide scanning. We thank Dr. Vishva M. Dixit (Genentech) for providing mutant mice. We thank Zsolt Czimmerer and Ana Rita Pombo Antunes for critically reading the manuscript. M.K. is supported by doctoral grants from Research Foundation Flanders (FWO, 1S23316N) and Kom op Tegen Kanker (Stand up to Cancer), the Flemish Cancer Society. E.L. is supported by a doctoral grant from FWO (1S67419N). A.M. is supported by a doctoral grant from FWO (1S16718N). H.V.D. is supported by a doctoral grant from FWO (1S24117N). S.P. is supported by a doctoral grant from FWO (1S68420N). D.L. is supported by grants from FWO (12Z1820N), Kom op Tegen Kanker and Vrije Universiteit Brussel. A.W. and J.V.G. are supported by Kom op Tegen Kanker (STIVLK2017000401). A.W. is supported by FWO (3G.0447.18).

## AUTHOR CONTRIBUTIONS

M.K. designed and performed experiments, analyzed data and wrote the manuscript. L.V.W., E.L., H.V.D., A.M., M.E., S.P., E.B., J.K., Y.E., M.S.M. and A.F. performed experiments and analyzed data. J.Q. performed computational analyses. D. Lambrechts, M.M., and A.W. acquired funding support and supervised data analysis. M.L., J.V.G. and D. Laoui acquired funding support, supervised the study and edited the manuscript.

