## Supplementary Figures for "Inflammasome- and gasdermin D-independent IL-1β production mobilizes neutrophils to inhibit antitumor immunity"

Supplementary Figure 1.

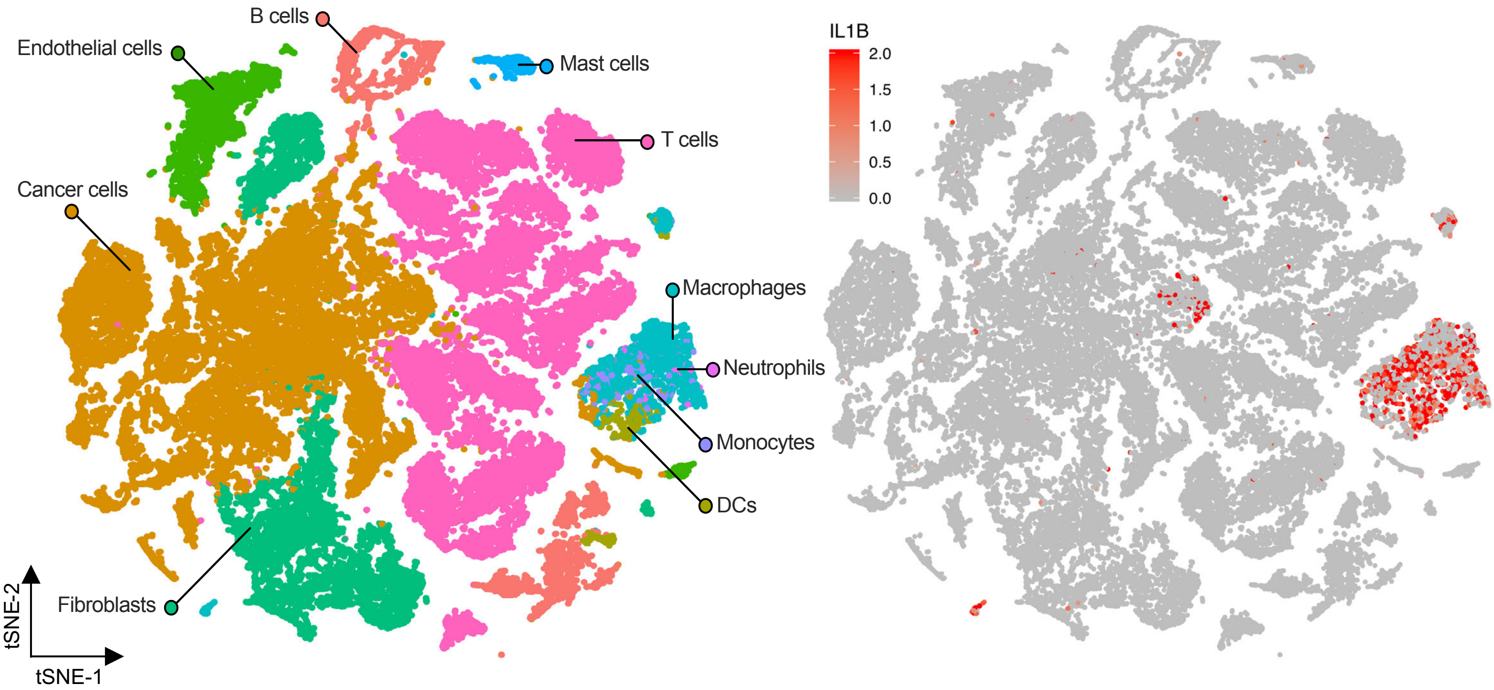

**Supplementary Figure 1.**
tSNE plot of cells from pooled human breast tumors (n=14) color-coded based on cell clusters or *IL1B* expression level.

Supplementary Figure 2.

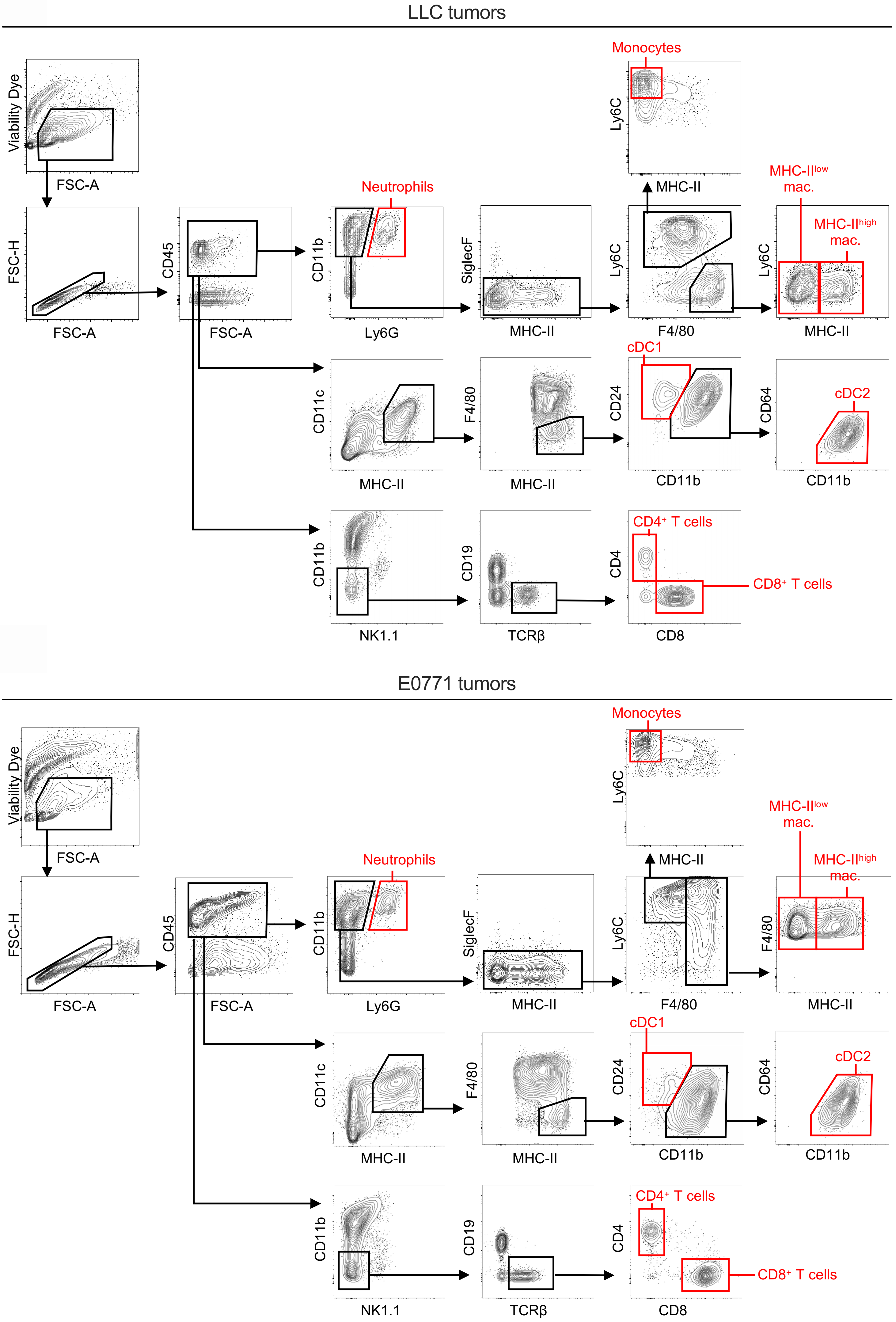

**Supplementary Figure 2.**

Representative flow cytometry gating strategies to identify immune cell populations in LLC and E0771 tumors used throughout the study.

Supplementary Figure 3.

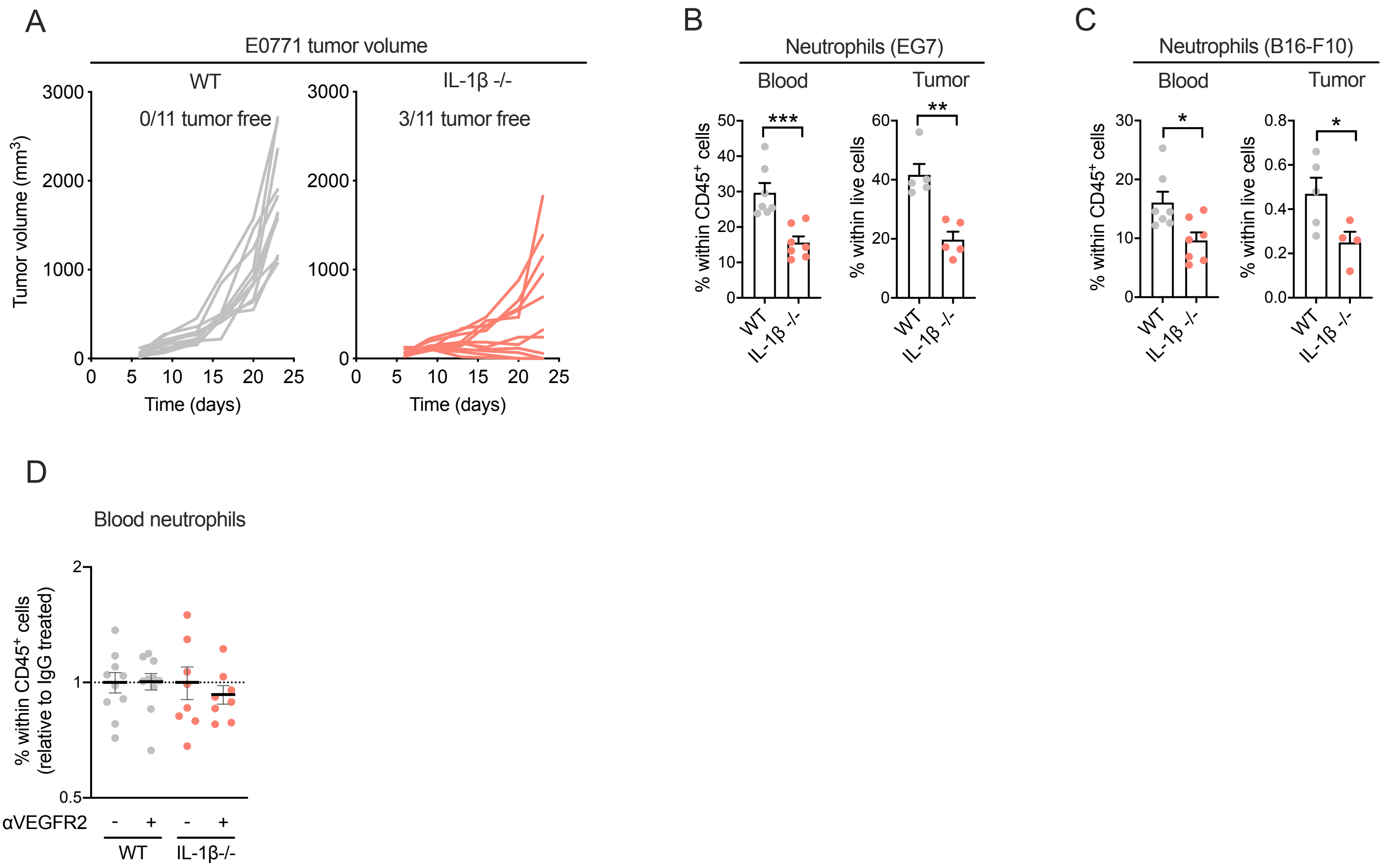

**Supplementary Figure 3.**
(A) Tumor volumes of individual mice with E0771 tumors from experiment shown in Figure 2A.
(B) Frequency of neutrophils in the blood (n=7) and tumor (n=5) of WT and IL-1β-/- mice with EG7 tumors.
(C) Frequency of neutrophils in the blood (n=7) and tumor (n=4-5) of WT and IL-1β-/- mice with B16-F10 tumors.
(D) Frequency of blood neutrophils in LLC tumor-bearing WT and IL-1β-/- mice treated with anti-VEGFR2 or IgG isotype control antibody normalized to the IgG isotype-treated control group within each genotype (n=7-10).
Graphs show mean and SEM.

Supplementary Figure 4.

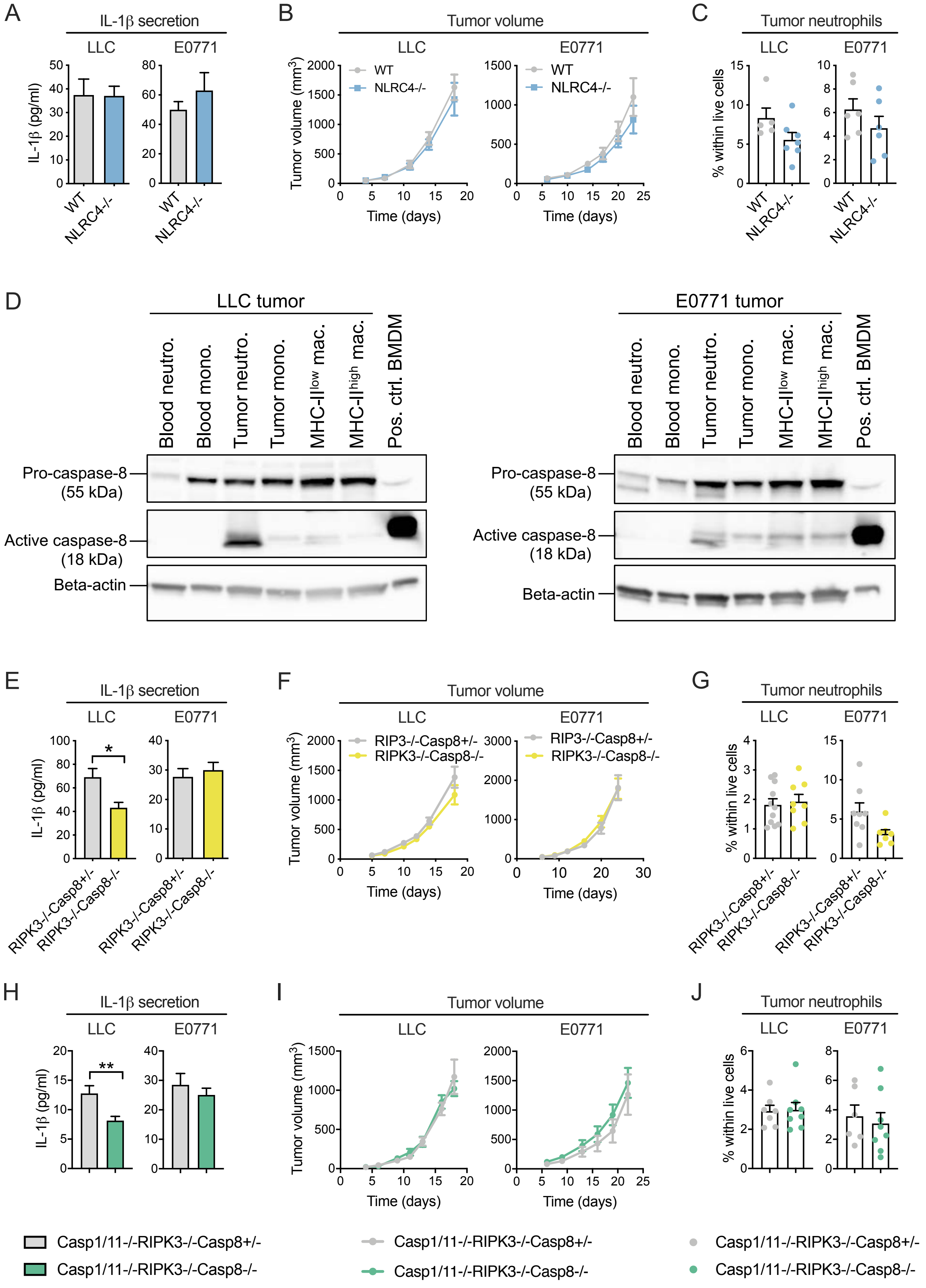

**Supplementary Figure 4.**

(A) IL-1β secretion of the $CD11b^+$ fraction of LLC (n=5-7) and E0771 tumors (n=6) from WT and NLRC4-/- mice measured by ELISA following 24 h *in vitro* culture.

(B) LLC (n=5-6) and E0771 (n=6-7) tumor growth in WT and NLRC4-/- mice.

(C) Frequency of neutrophils in LLC (n=5-7) and E0771 (n=6) tumors in WT and NLRC4-/- mice

(D) Immunoblots of caspase-8 on cell extracts of monocytes, macrophages and neutrophils isolated from LLC and E0771 tumors as well as monocytes and neutrophils isolated from the blood of tumor-bearing mice. Mouse bone-marrow derived macrophages (BMDM) treated with anthrax lethal toxin for 2 h were used as positive controls.

(E) IL-1β secretion of the $CD11b^+$ fraction of LLC (n=7) and E0771 tumors (n=7) from RIPK3-/-Casp8+/- and RIPK3-/-Casp8-/- mice measured by ELISA following 24 h *in vitro* culture.

(F) LLC (n=8-11) and E0771 (n=7-9) tumor growth in RIPK3-/-Casp8+/- and RIPK3-/-Casp8-/- mice.

(G) Frequency of neutrophils in LLC (n=8-11) and E0771 (n=7-8) tumors in RIPK3-/-Casp8+/- and RIPK3-/-Casp8-/- mice.

(H) IL-1β secretion of the $CD11b^+$ fraction of LLC (n=7-8) and E0771 tumors (n=6-8) from Casp1/11-/-RIPK3-/-Casp8+/- and Casp1/11-/-RIPK3-/-Casp8-/- mice measured by ELISA following 24 h *in vitro* culture.

(I) LLC (n=6-9) and E0771 (n=6-8) tumor growth in Casp1/11-/-RIPK3-/-Casp8+/- and Casp1/11-/-RIPK3-/-Casp8-/- mice.

(J) Frequency of neutrophils in LLC (n=7-8) and E0771 (n=6-8) tumors in Casp1/11-/-RIPK3-/-Casp8+/- and Casp1/11-/-RIPK3-/-Casp8-/- mice.

Neutrophil frequencies were determined by flow cytometry. Graphs show mean and SEM.

Supplementary Figure 5.

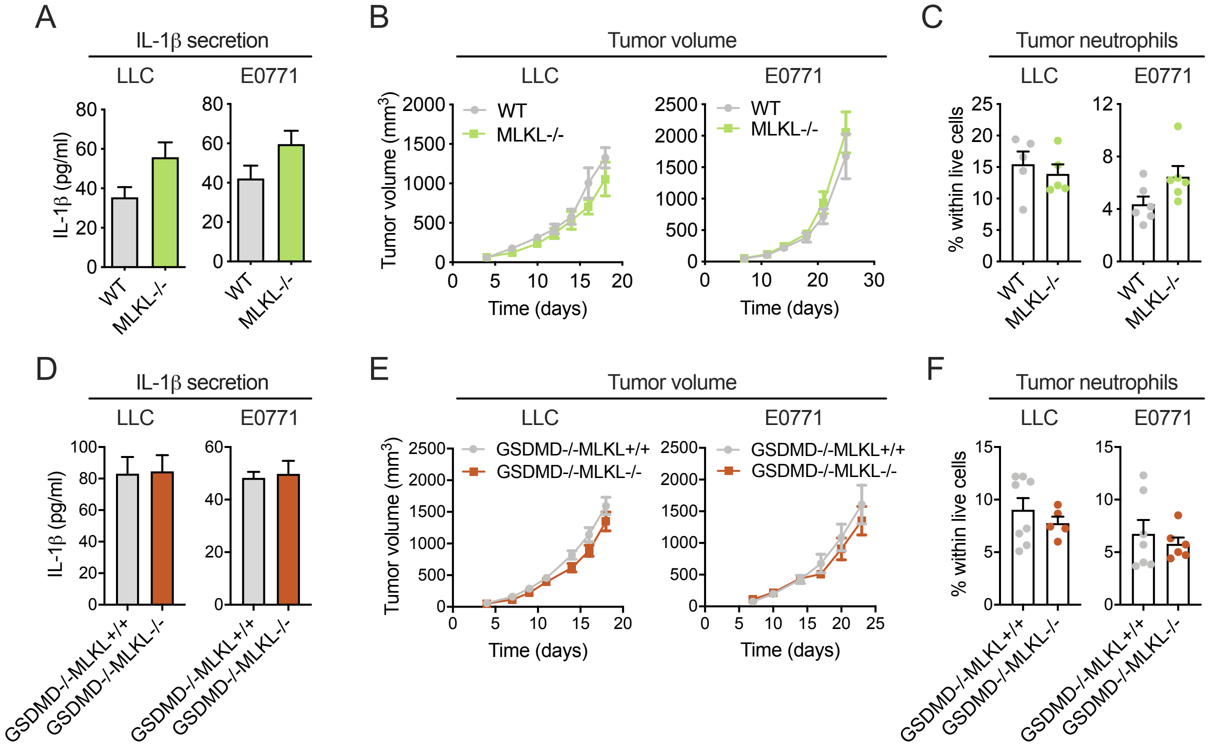

**Supplementary Figure 5.**

(A) IL-1β secretion of the $CD11b^+$ fraction of LLC (n=5) and E0771 tumors (n=5-6) from WT and MLKL-/- mice measured by ELISA following 24 h *in vitro* culture.

(B) LLC (n=5) and E0771 (n=9-12) tumor growth in WT and MLKL-/- mice.

(C) Frequency of neutrophils in LLC (n=5) and E0771 (n=6) tumors in WT and MLKL-/- mice.

(D) IL-1β secretion of the $CD11b^+$ fraction of LLC (n=7-8) and E0771 tumors (n=6-7) from GSDMD-/-MLKL+/+ and GSDMD-/-MLKL-/- mice measured by ELISA following 24 h *in vitro* culture.

(E) LLC (n=5-8) and E0771 (n=6-7) tumor growth in GSDMD-/-MLKL+/+ and GSDMD-/-MLKL-/- mice.

(F) Frequency of neutrophils in LLC (n=5-8) and E0771 (n=6-7) tumors in GSDMD-/-MLKL+/+ and GSDMD-/-MLKL-/- mice.

Neutrophil frequencies were determined by flow cytometry. Graphs show mean and SEM.

Supplementary Figure 6.

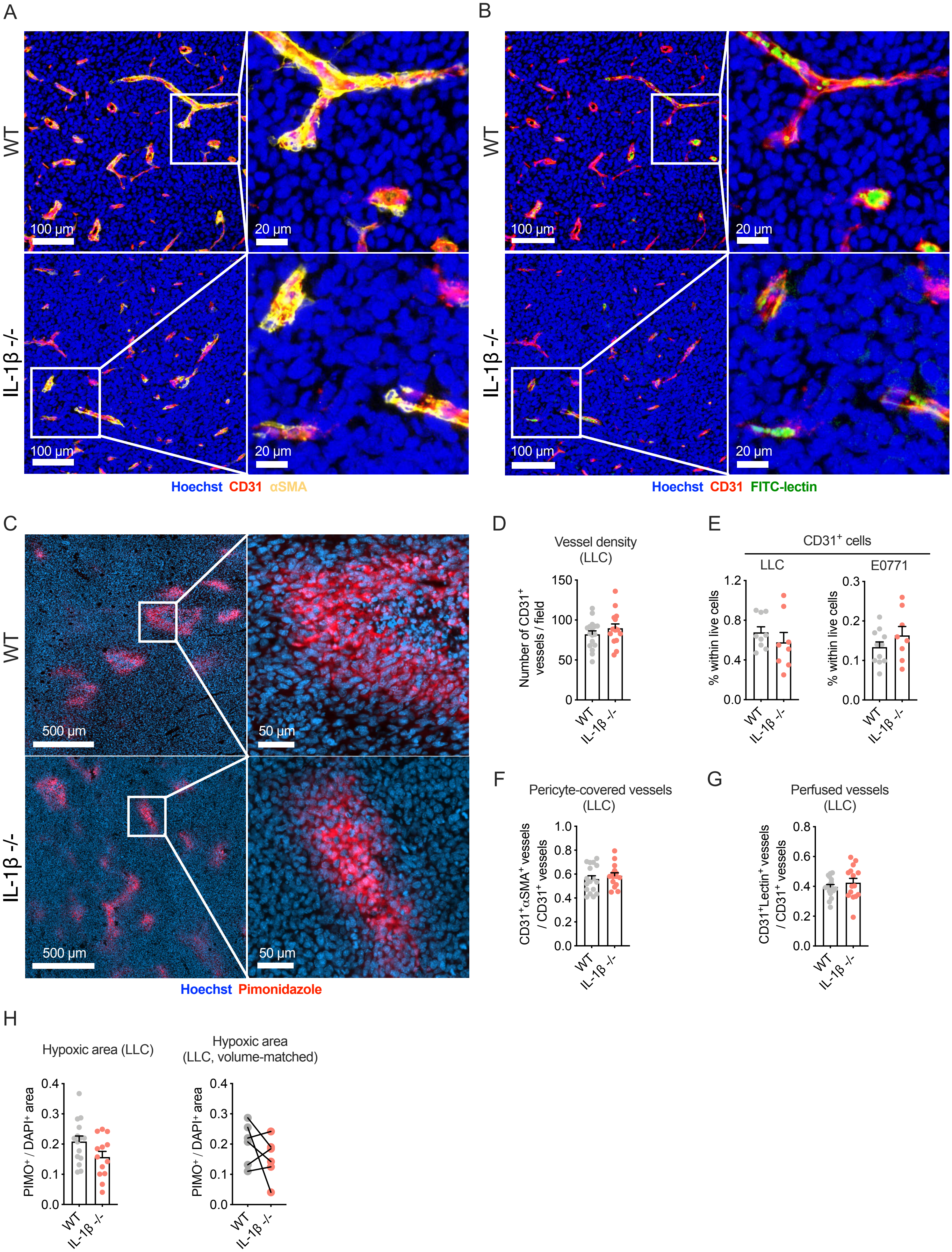

**Supplementary Figure 6.**

(A)-(C) Representative microscopic images of LLC tumor sections with immunofluorescent staining for CD31 (endothelial cells), αSMA (pericytes), FITC-lectin (perfused vessels) and pimonidazole (hypoxia) from WT and IL-1β -/- mice.

(D) Average density of $CD31^+$ blood vessels per high power field in LLC tumors of WT and IL-1β -/- mice (n=15-17).

(E) Frequency of $CD31^+$ endothelial cells in LLC (n=8-9) and E0771 (n=8-11) tumors of WT and IL-1β -/- mice quantified by flow cytometry.

(F) Proportion of $\alpha SMA^+$ pericyte-covered blood vessels in LLC tumors of WT and IL-1β -/- mice (n=15-17).

(G) Proportion of FITC-$lectin^+$ perfused blood vessels in LLC tumors of WT and IL-1β -/- mice (n=15-17).

(H) Proportion of $pimonidazole^+$ hypoxic areas in LLC tumor cross sections from WT and IL-1β -/- mice (n=13-15) and in volume-matched WT-IL-1β -/- tumor pairs.

Bar graphs show mean and SEM.

Supplementary Figure 7.

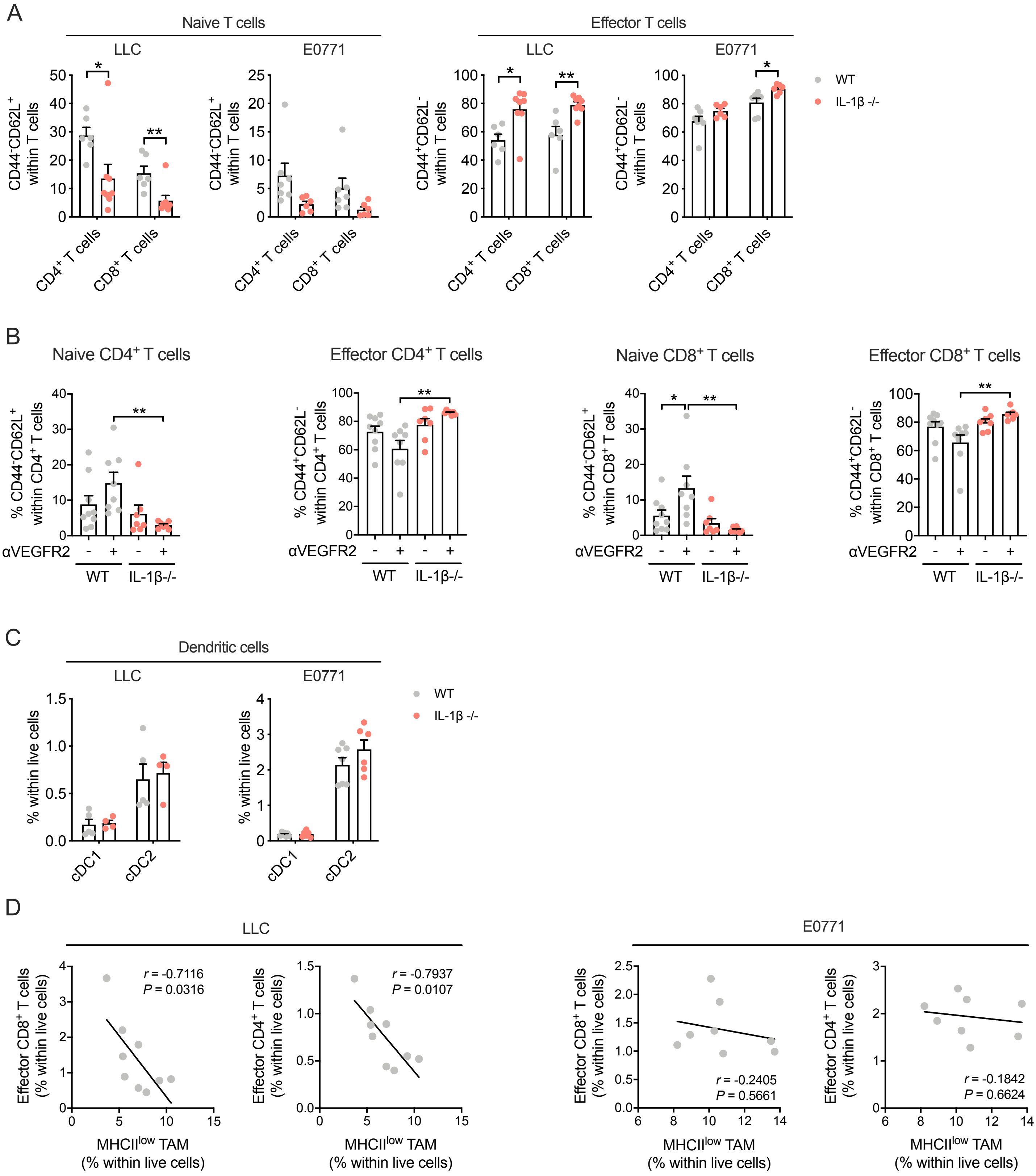

**Supplementary Figure 7.**

(A) Frequency of naive ($CD44^{-}CD62L^{+}$) and effector ($CD44^{+}CD62L^{-}$) T cells in LLC (n=6-8) and E0771 (n=6-7) tumors of WT and IL-1β -/- mice quantified by flow cytometry.

(B) Frequency of naive ($CD44^{-}CD62L^{+}$) and effector ($CD44^{+}CD62L^{-}$) T cells determined by flow cytometry in LLC tumors treated with anti-VEGFR2 or isotype control antibody (n=7-9).

(C) Frequency of type 1 and type 2 conventional dendritic cells (cDC) in LLC (n=4-5) and E0771 (n=6-7) tumors of WT and IL-1β -/- mice quantified by flow cytometry.

(D) Correlation of $CD44^{+}CD62L^{-}$ effector T cell and MHC-$II^{low}$ TAM abundance in LLC (n=9) and E0771 (n=8) tumors. Pearson *r* values and *P* values are indicated in the graphs.

Graphs show mean and SEM.

Supplementary Figure 8.

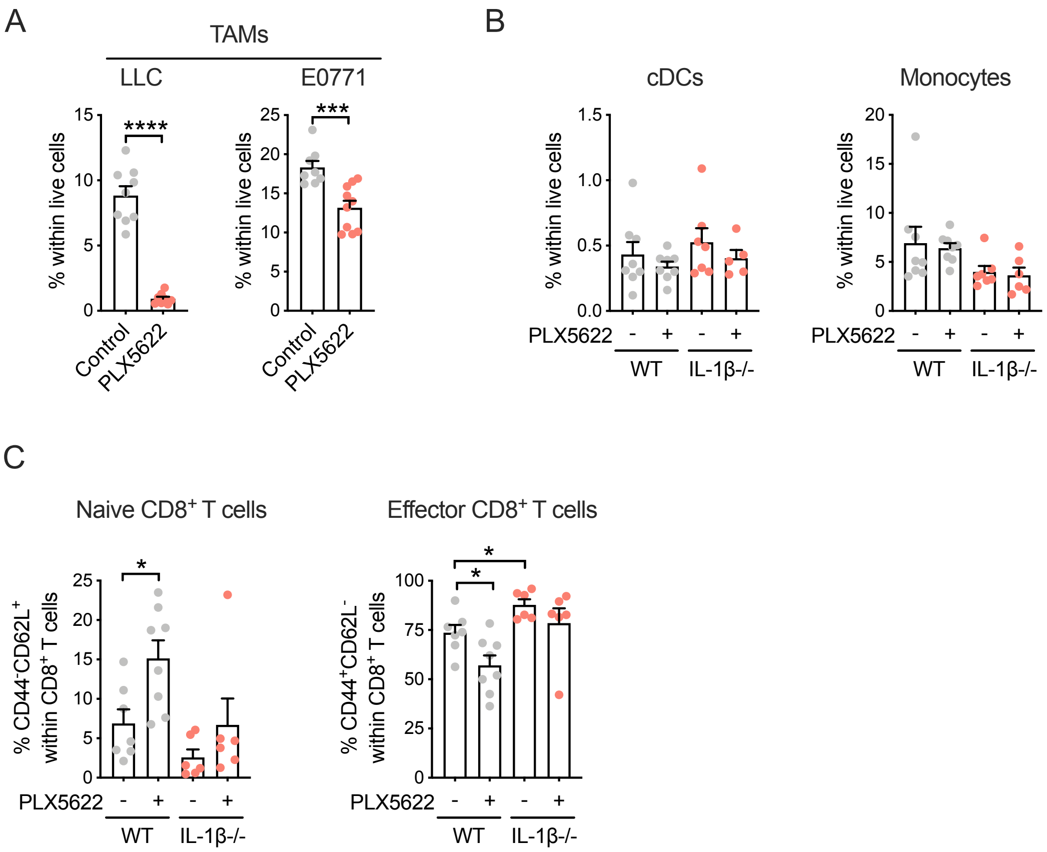

**Supplementary Figure 8.**

(A) Frequency of tumor-associated macrophages (TAM) in LLC (n=8-9) and E0771 (n=8-10) tumors in WT mice treated with PLX5622 or control diet quantified by flow cytometry.

(B) Frequency of conventional dendritic cells (cDC) and tumor-infiltrating monocytes in LLC tumors of WT and IL-1β-/- mice treated with PLX5622 or control diet quantified by flow cytometry (n=5-8).

(C) Frequency of naive ($CD44^{-}CD62L^{+}$) and effector ($CD44^{+}CD62L^{-}$) T cells determined by flow cytometry in LLC tumors of WT and IL-1β-/- mice treated with PLX5622 or control diet quantified by flow cytometry (n=6-8).

Graphs show mean and SEM.
